# Identifying the diamond in the rough: a study of allelic diversity underlying flowering time adaptation in maize landraces

**DOI:** 10.1101/092528

**Authors:** J. Alberto Romero-Navarro, Martha Wilcox, Juan Burgueño, Cinta Romay, Kelly Swarts, Samuel Trachsel, Ernesto Preciado, Arturo Terron, Humberto Vallejo Delgado, Victor Vidal, Alejandro Ortega, Armando Espinoza Banda, Noel Orlando Gómez Montiel, Ivan Ortiz-Monasterio, Félix San Vicente, Armando Guadarrama Espinoza, Gary Atlin, Peter Wenzl, Sarah Hearne, Edward Buckler

## Abstract

Landraces (traditional varieties) of crop species are a reservoir of useful genetic diversity, yet remain untapped due to the genetic linkage between the few useful alleles with hundreds of undesirable alleles^1^. We integrated two approaches to characterize the genetic diversity of over 3000 maize landraces from across the Americas. First, we mapped the genomic regions controlling latitudinal and altitudinal adaptation, identifying 1498 genes. Second, we developed and used F-One Association Mapping (FOAM) to directly map genes controlling flowering time across 22 environments, identifying 1,005 genes. In total 65% of the SNPs associated with altitude were also associated with flowering time. In particular, we observed many of the significant SNPs were contained in large structural variants (inversions, centromeres, and pericentromeric regions): 29.4% for flowering time, 58.4% for altitude and 13.1% for latitude. The combined mapping results indicate that while floral regulatory network genes contribute substantially to field variation, over 90% of contributing genes likely have indirect effects. Our strategy can be used to harness the diversity of maize and other plant and animal species.

---

Maize *(Zea mays* subsp. *mays*) is a model organism with a legacy of a hundred years of cytological, genetic, and biomolecular characterization^2^. Maize displays high levels of genetic diversity with low linkage disequilibrium (LD)^3,4^, low population differentiation^5^, prevalent migration^6^ and occasional introgression from wild relatives^7–9^. More recently, experimental populations like the Nested Association Mapping (NAM) populations ^10,11^, and large association panels^4,12^ have allowed mapping and deployment of useful alleles for several quantitative traits^13–16^. However, most of the founder lines from these panels correspond to highly inbred improved lines, many from temperate regions, capturing only a modest fraction of the total diversity present in the species. In contrast, maize landraces span numerous ecogeographic areas and harbor most of the diversity of the species. Nevertheless, maize landraces like many other crops traditional varieties remain largely uncharacterized by genomics.

This study maps genes controlling flowering with two distinct methods: (1) Each of these landraces come from environments to which they are well adapted. We used this adaptation as the trait to identify genes driving large scale adaptation. (2) We mapped flowering time variation in controlled field experiments through a novel, rapid, experimental design called F-One Association Mapping (FOAM) (Figure 1). Briefly, FOAM consists of sampling single individuals across numerous populations, which are genotyped and crossed to one or a small number of common parents to derive F1 families. Subsequently GWAS is performed from multi-trial F1 progeny evaluation. Major advantages for this design are (a) capturing thousands of alleles across populations, (b) maintaining the tractability of two alleles per loci per individual, (c) ample replication of alleles increasing the power and accuracy for genetic effect estimation. The main limitation of FOAM is that the nested evaluation of different subsets of F1 progeny by ecological zone limit the ability to accurately estimate genotype by environment interaction effects.

**Figure 1.**
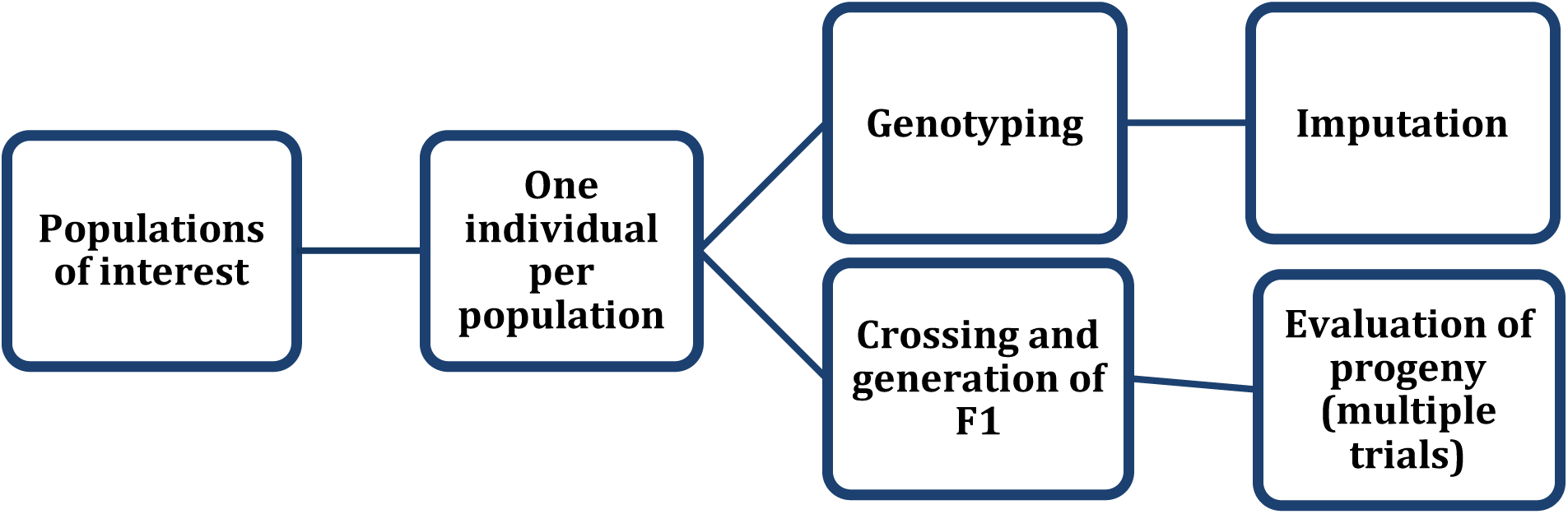
Experimental design. One individual from each of up to thousands of individuals is genotyped and used as parent. Progeny are then evaluated for multiple years/locations to estimate the genetic contribution of the original individual and phenotypic and genotypic data are used for Genome Wide Association

Our maize landrace FOAM population used individuals from 4,471 accessions from 35 countries in the Americas (Figure 2) grouped into three adaptation classes to account for altitude adaptation (low, middle and high elevation). Similarly, the common parents and evaluation sites were nested within adaptation class (methods, supplemental figure 1) ^17,18^. Landrace parents were genotyped for close to one million SNPs using Genotyping by Sequencing^19^, and missing data was imputed using BEAGLE4^20^. Of the 4,471 accessions, 3,552 yielded F1 families containing both genotypic profiles and sufficient progeny, 3,633 contained detailed passport information which was used for mapping large scale adaptation, and 2,603 were present in both mapping studies.

**Figure 2.**
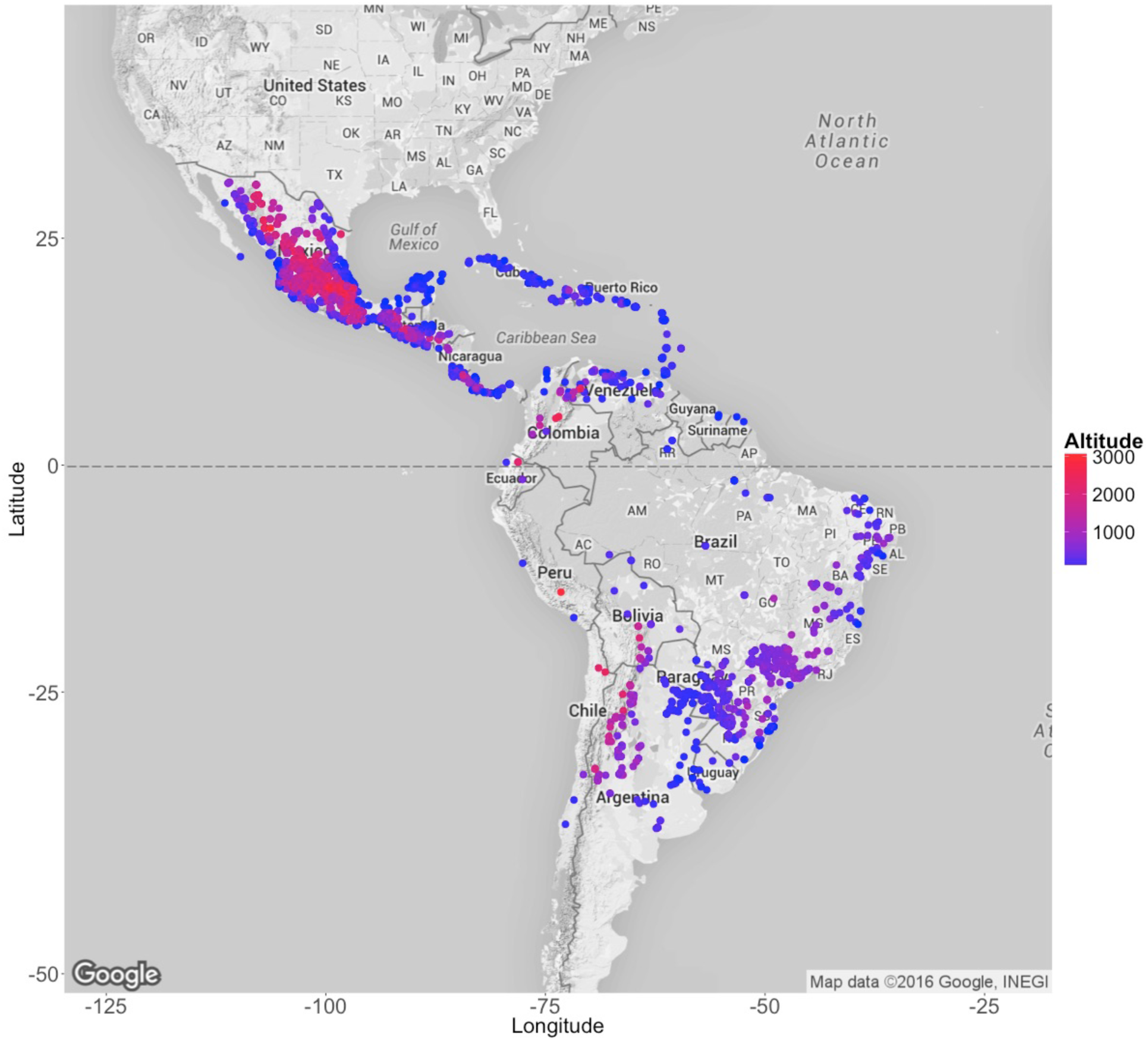
Geographic coordinates of original sampling sites of landrace accessions. Color gradient corresponds to altitude

We first explored the effects of recombination frequency and geography-driven limited dispersal on the distribution of genetic diversity in the landrace parents. Using Multidimensional Scaling (MDS, Methods), we observed the first axis and second axes explained only 6.1% and 1.7% of of the variance respectively, consistent with the low F_ST_ in maize landraces^5^. The first axis separates among Mexican landraces, consistent with Mexican landraces having a deeper coalescent and greater representation in the panel. The second axis was associated to a latitudinal North to South gradient across Latin America representing isolation by distance (Supplemental Figure 2). In addition, a Mantel test^21^ revealed a significant correlation between geographic and genetic distances (Pearson's r= 0.46, P-value<.001), with most of the association driven by altitude. Despite this, phylogenetic analysis (Methods, Supplemental Figure 3) revealed that adaptation class does not drive clade membership, which indicates that alleles segregate across adaptation classes, with highland adaption being polyphyletic, consistent with recent reports^22^. To study recombination, we estimated an approximate LD statistic (Methods) which shows a distribution consistent with previous recombination estimates^23,24^, with higher recombination in gene-rich regions, and lower around centromeres. Each chromosome displayed a unique recombination landscape, with the presence of half a dozen high LD regions (Supplemental figure 4), which together encompassed 6.1% of the base pairs of genome, accounting for 2.8% of the annotated coding genes. Together, these results suggest that although at large scale geography and adaptation contribute to the distribution of diversity, even with the large effective population size of landraces at the genomic scale a complex recombination landscape limits the free segregation of alleles through increased LD.

Flowering time generally plays a crucial role in local adaptation of plants, and in maize flowering time is a complex trait controlled by hundreds of small effect loci, many with rich allelic series ^4,14,25–30^. We used altitude and latitude from sampling location as traits for mapping local adaptation, and the significance thresholds were chosen to maximize genic overlap rate between flowering, altitude, and latitude (Methods, supplemental figure 5). For altitude, we observed 58.4% of the significant SNPs corresponded to regions with higher LD. In particular, INV4m, the 13Mb adaptive introgression from highland teosinte into maize^8,31^ was highly significant. We also observed significance for the centromeres of chromosomes 2,5,6,8 and a large region upstream of the centromere on chromosome 3. Outside this low recombination regions, 366 genes showed significant association with altitude. For Latitude, we observed 13.1% of the significant markers were contained within low recombination regions, particularly the centromere of chromosomes 5. In total across all Latin America, 1498 genes showed significant association with latitude, of which 395 of were shared with altitude. The minor allele frequency distribution of the significant alleles indicated that many are shared across clades and landraces, which was very distinct from the neutral distribution (Figure 3). These 1498 genes appear to be the main contributor to large scale environmental adaptation to altitude and latitude — the key drivers of flowering time.

**Figure 3.**
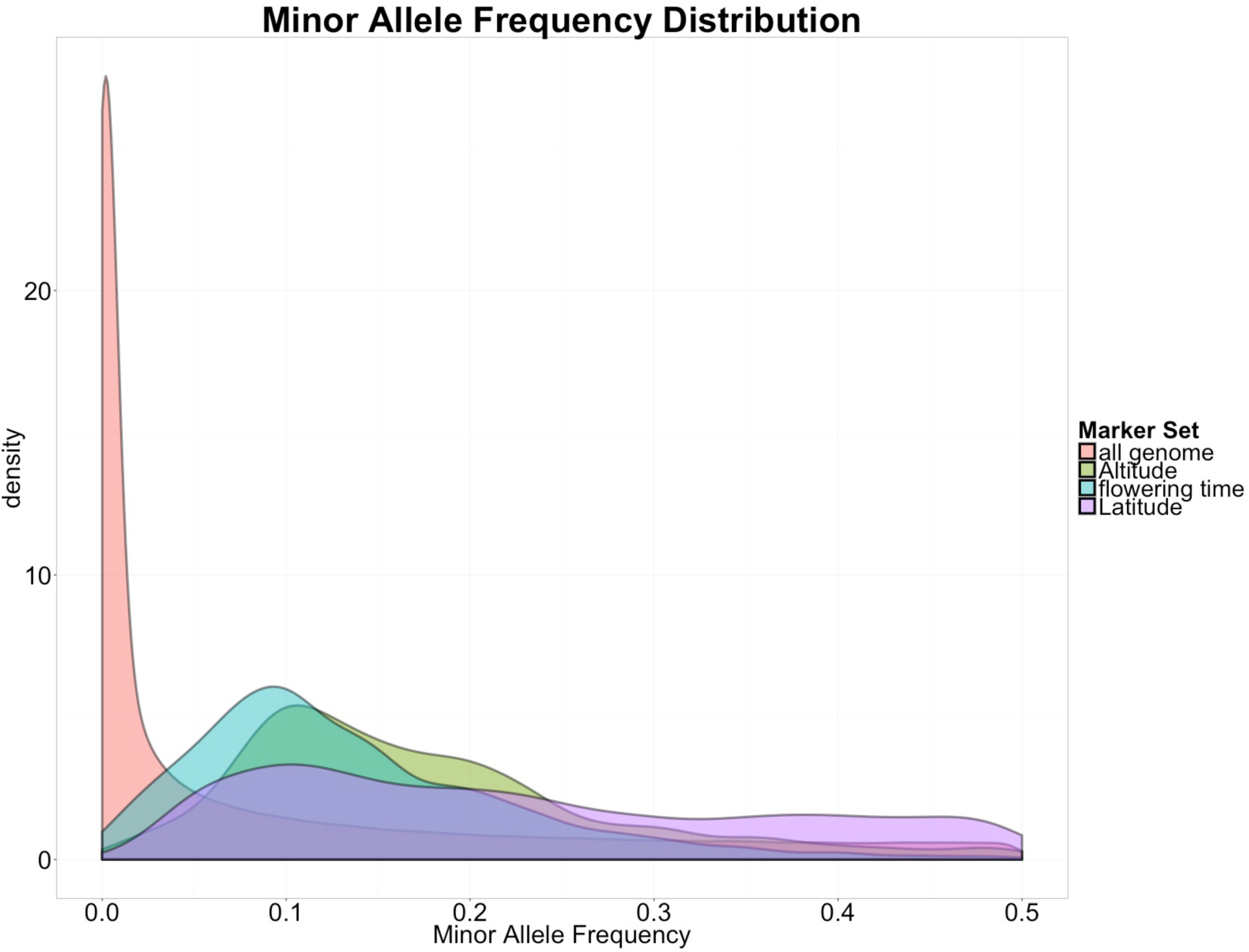
Minor Allele frequency distribution for all segregating SNPs, as well as the most significant SNPs for each trait.

To study the genetic basis of flowering time, we conducted field evaluation on F1 progeny across 22 trials and 2 years in 13 locations across Mexico, with each trial containing a different subset of the collection to maximize number of accessions evaluated (Methods, supplemental table 1). Phenotypic data was analyzed independently for each trial using a mixed linear (Methods), yielding 18,797 accession parent-environment estimates for each male and female flowering time. We performed genome wide association for days to male and female flowering using a mixed linear model (Methods). In total 72% of the associated SNPs were significant for both male and female flowering, as expected from the overlapping genetic control^14^. There was a significant contribution of low recombination regions in flowering time variation, parallel to that of latitude and longitude, with a 20-fold enrichment for significant SNPs at high LD regions (Pearson's chi-squared, p-value < 2.2e-16). In particular, significant variants included the centromeres of chromosomes 3, 5, and 6, INV4m, and a 6Mb region on chromosome 3 beginning at 79Mb. The 6Mb region on chromosome 3 has a segregation similar to INV4, and together with its increased LD suggests it might be an inversion. In NAM this putative inversion and the centromere comprise a single QTL for flowering time^14^. For the centromere of chromosome 5 there were 3 distinctive alleles segregating in the landraces, all present in the NAM population (supplemental figure 6). The inverted allele of INV4m, although absent in temperate material, segregates at high frequency in highland landraces (supplemental figure 7), where it has very large additive effect advancing flowering by three days, the largest effect for flowering time in maize to date. Both homozygous alleles from the putative inversion on chromosome 3 segregate across our maize landrace panel and the NAM population. Compared to INV4m, this locus displays a more modest effect on flowering time. The heterotic effect of the centromere of chromosome 5 on yield^32^,potentially product the complementation of deleterious mutations^23^, suggests that the significant inversions and centromeres may similarly affect flowering time through heterotic effects leading to more vigorous plants, which in maize generally results in earlier flowering.

Outside the structural variants, we observed 881 and 883 genes (around 2.2% of genes) with significant association for days to female and days to male flowering respectively (Supplemental Tables, Figure 4). To further characterize the regions associated with flowering time, we looked for gene ontology enrichment and gene expression using the maize transcription atlas^33^(Methods), and compared the significant genes to a candidate gene list containing genes characterized in other populations, known to interact in the maize flowering time regulatory network^34^ as well as the 25 members of the *Zea mays* CENTRORADIALIS (ZCN) gene family ^35^. Overall the associating genes tended to be expressed in anthers, and enriched in general metabolic processes, with the genes known to be part of the regulatory network being more expressed in immature cob and the tip of the leaf at V5 stage and enriched for regulatory processes (Supplemental figure 8). We observed a significant enrichment in flowering time candidate genes compared to the rest of the genome (Fisher's Exact Test p-value = 4.3 x 10^-7^). In total 10 and 12 candidate genes representing the circadian clock, photoperiod, gibberellin acid, and circadian clock pathways displayed significant associations with male and female flowering respectively. Out of these, nine were common for both types of flowering. The most significant hits corresponded to VGT1^36,37^, one of the largest known G×E QTL, and ZCN8^35,38^, the maize florigen and homolog to *FT* in *Arabidopsis* (Figure 5). ZmCCT, the largest photoperiod QTL^29^ was only modestly significant for latitude, and significant only for days to female flowering, probably a result of non-inducing sampling and trial locations. In particular for the gene *d8,* a locus with cryptic association with flowering time^34^, we observed significance for this gene around 50 and 100kb up and downstream the coding region for latitude, altitude, and both male and female flowering, overlapping with the region previously observed to display divergent selection associated with climate adaptation^39^. In addition, the distribution of the flowering time associating genes displayed a significant geography effect, with 56 and 52 genes in common with altitude and latitude respectively. In general, the minor alleles for flowering time tended to be associated with high elevation, and northwest coordinates, however the minor allele frequency distribution of the significant SNPs was different to that of the alleles significant for altitude and latitude, having a significant shift towards low frequency polymorphisms (Figure 3). Together, these results support the model of infrequent variants in recurrent regulatory genes underlying the genetic control of flowering time variation in maize, with adaptive alleles segregating across populations, and their distribution matching the fitness optimum according to geographic variation. In particular, the high overlap between significant SNPs for altitude and flowering time suggests that for tropical maize flowering time adaptation is very relevant for changes in elevation, which affects among others spectral composition and intensity of incident light, as well as the incidence of heat and cold stress. In contrast, the lower overlap between latitudinal and flowering time associating SNPs could be to the sampling from non-photoperiod inducing latitudes, potentially leading to latitudinal flowering time adaptation being relevant for other biotic (disease pressure) and abiotic (soil pH, precipitation) stresses.

**Figure 4.**
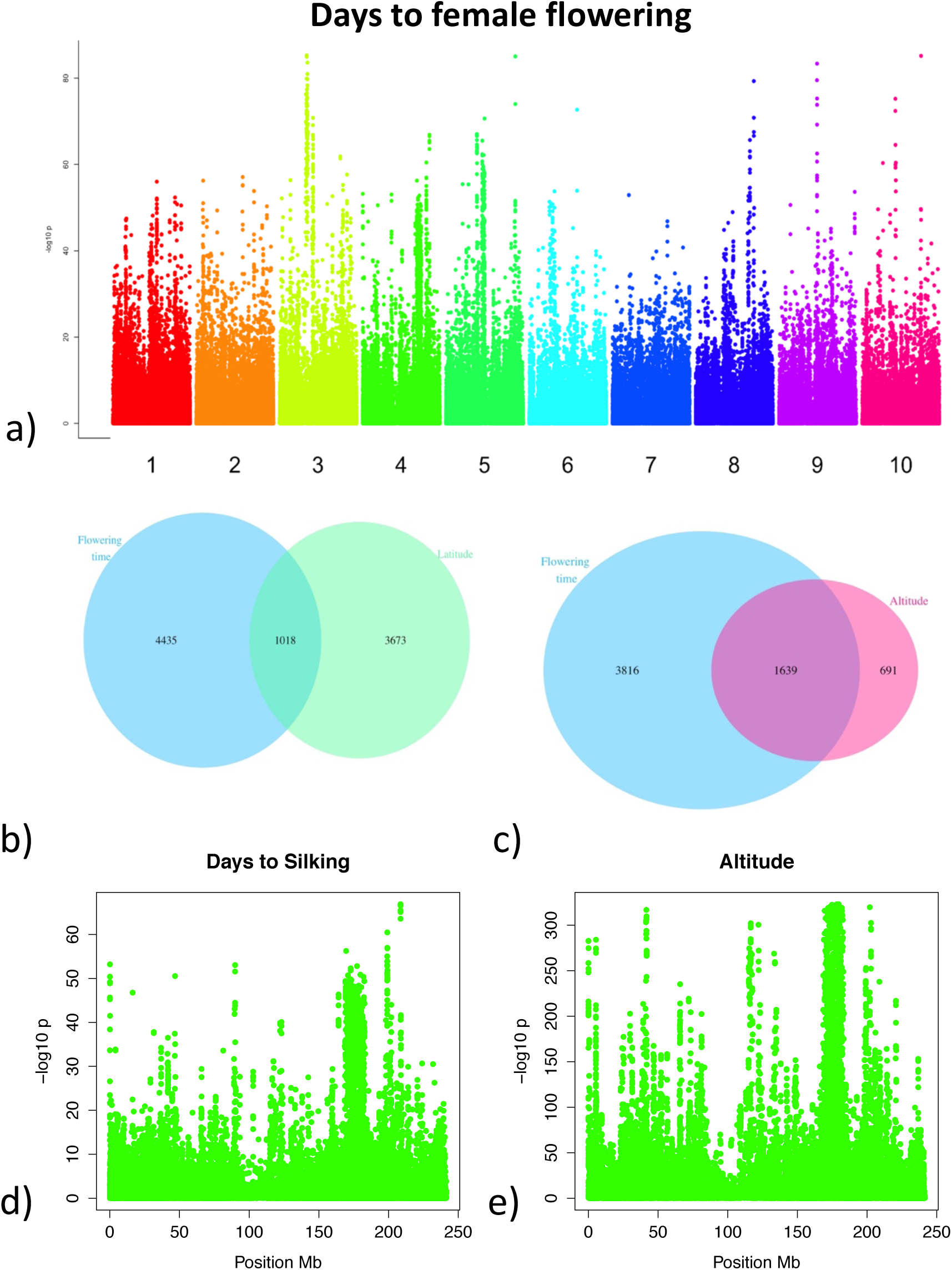
a) Manhattan plot for Days to silking. b) Local Manhattan plot for chromosome 4 for days to silking and c) Altitude. The large region with significance corresponds to INV4m, the adaptive introgression from highland teosinte to highland maize d) Overlap between significant SNPs for flowering time and latidue and e) altitude

**Figure 5.**
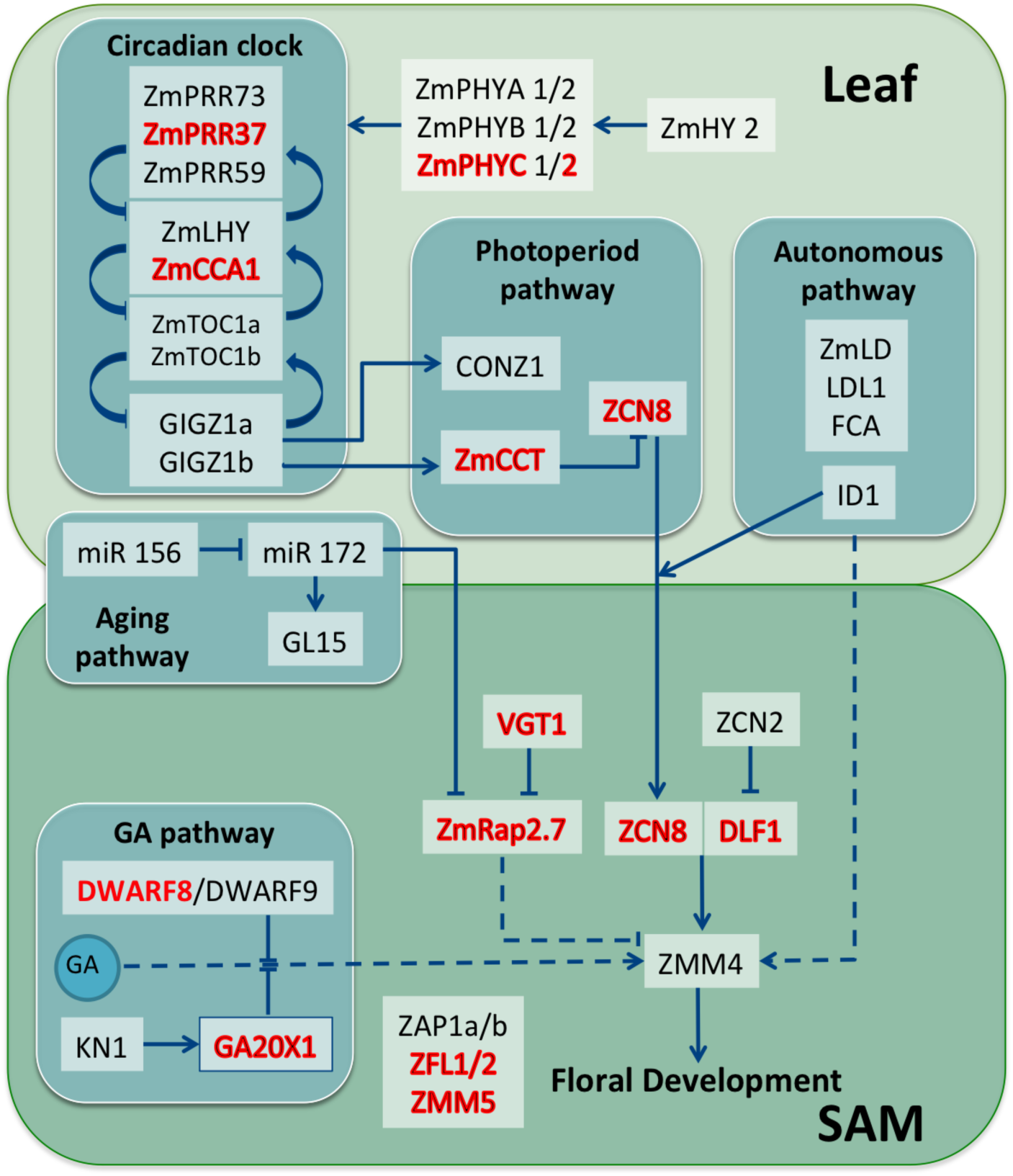
Flowering time pathway from Dong, *et al^34^,* showing the genes involved in flowering time at the leaf and Shoot Apical Meristem (SAM). The genes highlighted in red displayed significant association with flowering time in our study.

We assayed the potential for predicting flowering time in the landraces using either all our high density genetic markers or just the markers significantly associated with the trait. We performed genome wide prediction using gBLUP independently for each trial (Methods). The average 5fold cross-validated prediction accuracy was 0.45 across trials for both male and female flowering time and, and as high as 0.7 for some trials (Supplemental figure 9). Genomic prediction accuracy between the top genes from GWAS was equivalent to that of 30,000 random evenly distributed SNPs, highlighting their potential use for breeding of the significant markers. Intriguingly prediction accuracy was not correlated with our other heritability estimate (Pearson cor=0.22), which could be an effect of the differences in the genetic variances and sample sizes across all trials. Together the good predictive ability of the significant regions for genomic selection shows the potential to greatly speed the breeding of new adapted varieties with exotic beneficial alleles.

Crop landraces are an incredible source of diversity that will be necessary to adapt our crops to next century of climate change. However, their tremendous diversity and genetic load prevent them from being efficiently tapped without a genomic index. This research lays out two complementary strategies for tapping this diversity. The geographic associations are powerfully identifying the adaptive loci, which appear to be common and shared, and are unlikely to be deleterious given their high frequency. This extensive sharing is probably the result of outcrossing and extensive migration throughout Latin America in last several millennia. The limitation of this approach is that correlated traits and adaptations are being co-mapped. The novel FOAM field trial associations, while substantially overlapping, are showing the impacts of deleterious and private mutations and their complementation in these hybrid trials. These deleterious alleles have been the bane of breeders wanting to tap landrace diversity. The strategy for tapping this diversity should be use the overlapping genes and alleles of the two separate approaches— as these have proven to be adaptive *and* target the trait of interest. The breeding could use standard genomic selection or genome editing. This provides an efficient strategy to tap landraces diversity and allow our crops to adapt to faster changes than ever had in the past.

## Methods

### Mating design and phenotypic evaluation

The mating design for the maize landrace FOAM population consisted of crossing each accession male to single cross hybrid females of matching adaptation. Leaf tissue of the landrace individual was collected for genotyping. The progeny evaluation trials were performed across 2 years in 13 locations across Mexico using an augmented row-column design, which includes systematic checks in field rows and columns^40^. There were between 288 and 1928 accessions per trial, with an average of 834 (Supplemental table 1). Over half of the accessions were replicated in 5 or more trials, with a maximum value of 13 trials per accession and a minimum of 1. For each trial, each experimental row contained between 10 and 25 progeny plants. The replication across trials together with the use of systematic checks across experimental fields provides sufficient allelic replication for accurate estimation of genetic effects. Flowering time was measured in each trial following the maize standard, i.e. the number of days from planting until half of the individuals within a plot displayed silks for female flowering or anthers in half of the central spike for male flowering.

### Genotyping

Accessions used as male parents were genotyped using GBS^19^, with ApeKI as the restriction enzyme to a replication level of ~96 individuals per sequencing plate. Approximately 8×10^9^ sequencing reads were generated using an Illumina HiSeq for the landrace accessions and sequence reads were analyzed jointly with another 40,000 maize lines as part of the GBS Build 2.7 using TASSEL^41^. For association analyses, missing data was imputed using BEAGLE4^20^, which has been shown to yield the best current accuracies in maize heterozygous material (R^2^=0.68)^42^. After imputation, SNPs were filtered for minor allele frequency greater than 1% resulting in approximately 500,000 biallelic markers across the genome. GBS non-imputed markers can be accessed at http://hdl.handle.net/11529/10034 and imputed GBS markers at http://hdl.handle.net/11529/10035

### Diversity Assessment

For the Mantel test^21^, we calculated the pairwise Euclidean distance matrix based on the geographical data from the accessions (latitude, longitude, and altitude, http://mgb.cimmyt.org/gringlobal/search.aspx). The genetic distance matrix was estimated from a genome wide random sample 30,000 non-imputed markers using TASSEL. The distance matrix was used for estimating the Neighbor-Joining tree using TASSEL. Multidimensional Scaling (MDS) was performed on the genetic distance matrix using the cmds function in R.

### Recombination

Our LD statistic consisted in estimating the correlation between markers across the genome at 100 site windows using all homozygote and heterozygote non-imputed markers with the LD function on the software TASSEL. For comparing the LD and recombination values, we estimated the correlation at 1Mb sliding windows between (1) the log10 median LD estimate (2) the log median crossover probabilities estimated using the American and Chinese Nested Association Mapping populations^23^, and (3) the log median population recombination rates (rho) estimated both for improved lines and landraces Hapmap2 project^24^. Our LD estimates displayed a negative correlation with gene density (r=-0.57) and NAM crossover probability^23^ (r=-0.45). We observed a modest negative correlation (r=-0.33) with a population genetic estimate of historical recombination (rho) ^23,24^. High-LD regions were defined based on the change in slope of global median LD (Figure 5- Figure supplement 1) as those segments that had a median LD greater than 0.01. In total, there were 256 high LD regions encompassing 7.8% of the genome.

### Genome wide association with altitude and latitude

We performed Genome Wide Association using a generalized linear model with altitude and latitude as response variables and markers filtered at 1% frequency as explanatory variables. Altitude and Latitude were recorded during field sampling of the original accessions. In order to establish a significance threshold to avoid excess of false positives, we estimated the overlap rate using the most significant flowering time GWAS SNPs. Overlap Rate was defined as the set of overlapping SNPs between the top flowering time SNPs and either altitude or latitude, divided by the union of the sets across significance thresholds from 0.001 to 0.01. Significance thresholds chosen were 0.005 for altitude and 0.01 for altitude (Supplemental figure 4). Heritability estimates were 0.88 for altitude and 0.85 for latitude, estimated LDAK^43^ with a single Kinship matrix, estimated with all the Beagle4 imputed markers, and the matrix was estimated from the algorithm implemented in GCTA^44^.

### Analyses of structural variants

In order to infer the underlying haplotypes for the centromeres of chromosomes 3,5,6, as well as INV4 and the high-LD region on chromosome 3, we first estimated a genetic distance matrix for each locus using the non-imputed markers. The distance matrices were then analysed using multidimensional scaling. The centromere of chromosome 5 segregates in the landraces with three distinct homozygous haplotypes and their corresponding heterozygote pairs. The region around the centromere of chromosome 6 was 12 Mb in size, includes the centromere and a large pericentromeric region that expands out in both directions; it displayed a similar pattern to the centromere of chromosome 5 however distinct alleles were not called due to the excess of heterozygous individuals between the homozygous classes, probably reflecting recombinant haplotypes. The centromere of chromosome 3 displayed a more complex pattern of distance than the other two associating centromeres, likely due to the presence of more than three segregating haplotypes. For INV4, we observe two distinct alleles and the heterozygote. We observed the allele is fixed in many of CIMMYT improved lines (Table 3), including those used as parents for the highland test crosses in the present experiment.

### Analysis of phenotypic data

To estimate the breeding values of the landrace accession parent, for each trial a mixed linear model was fitted using restricted maximum likelihood method, in ASREML (V 3.0), using the progeny’s calendar days to male or female flowering as a response variable. Of the 23 trials planted, one was excluded because flowering time data was not collected according to protocol. The models included fixed effects for checks, tester, and hybrid and a random effect of accession in a complete nested model. Including in the model the random effect of row and column and using an autoregressive model of order 1 in row and columns controlled experimental noise product of field variation. All the random effects were considered independent one from each other. The model used can be expressed as follows:

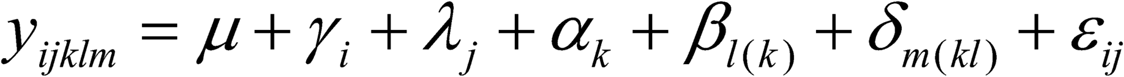

where

Y_ijklm_: is the repsonse variable,
μ: is the overall mean,
γ_i_: is the effect of the i-th row, 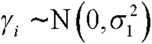
λ_j_: is the effect of the j-th column,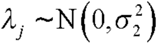
α^k^: is the effect of the k-th group, k=l,…,K,K+l, if k ≤ K the group is a check, the group K+l is the average of testers.
β_l(k)_: is the effect of the 1-th tester in group K+l
δ_m(kl)_: is the effect of the m-th accession in the tester k in group K+l,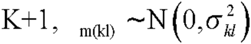
ε_ij_: is the experimental error

 for the experimental error we assume the following distribution:

ε ~N(0,Σ), with Σ=Σ_r_ ⊗ Σ_c_ and

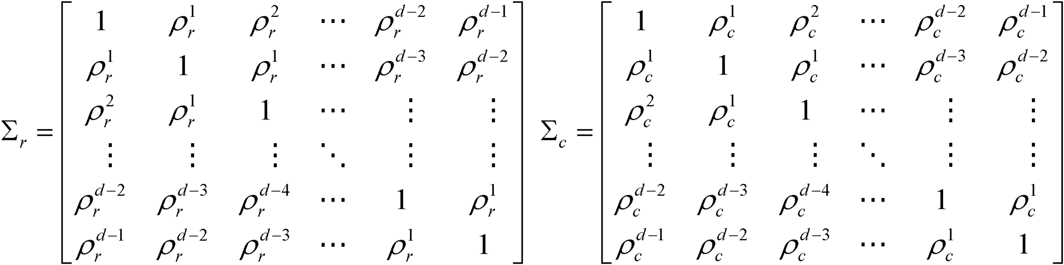

### Genome wide association with flowering time

Association analysis was performed in two steps for all trials using a linear mixed model. For each trait (days to male and female flowering) two models were fitted, one with the trait BLUPs as response variable and another one with the standardized values of the same BLUPs. This was done in the absence of growing degree units, to verify the consistency of the results given the uneven variances for the trait across the various trials. The first step models included the fixed effects for trial (categorical); population structure in the form of 10 MDS weights (numerical) that together explained around 13% of the genetic and 10.6% of the phenotypic variances; and the effect of the hybrid used as parent for each accession’s cross. The random effect of relatedness was added to both models in the form of a kinship matrix. The kinship matrix was estimated using the same subset of SNPs as the MDS weights. The mixed model was fit using the R package EMMREML (http://cran.r-project.org/web/packages/EMMREML/index.html). Residuals were obtained from those models and fitted in the second step models as a response variable for the single marker analysis using R, with marker nested within trial.

The model equation used was

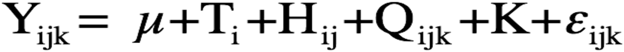

where

Y_ijklm_: is the repsonse variable,

*μ*: is the overall mean,

*T*_i_: is the effect of the i-th trial

*H*_j_ is the effect of the hybrid parent

*Q*_k_: is the population structure effect containing 10 weights from MDS

*K*: is the random effect of relatedness through kinship matrix K estimated from 30,000 random SNPS

*ε*_ijk_: is the residual error

In the second step of the association model, the residuals from the first model were fitted as a response variable in the following model

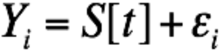

Where Y is the residual, S is the SNP effect and is nested within trial t. The model tests the null hypothesis that the effect of each SNP is 0 in all trials.

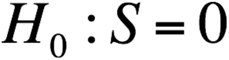

The alternative hypothesis is that the SNP has an effect on any trial. The reason for testing this hypothesis is that the effect of each SNP can, and often does, change on value and direction depending on its segregation on the population and its phase with the causal polymorphism. We consider as significant the top one percent of the SNPs based on p-value, which all had −log10 p-values greater than 18.

### Code availability

R implementation of the ASREML code used for estimation of breeding values can be found at http://data.cimmyt.org/dvn/dv/cimmytswdvn;jsessionid=c1de29cab7c37b41098fd8ad6684

The mixed model was fit using the R package EMMREML (http://cran.rproject.org/web/packages/EMMREML/index.html)

All other additional scripts are available through github user

### Significance at genic regions

We reasoned that significance at candidate genes would depend on local LD and genotype coverage, therefore a higher proportion of significant SNPs around candidate genes would be indicative of association at the gene itself rather than at the entire LD block or because of higher genotype coverage. On that account, we looked at significant associating SNPs within 50 kb up and downstream of candidate genes. Of all the candidate genes, only PhyB1, GL15 and ZCN13 are in the high-LD set and therefore were excluded from this analysis.

Genome wide prediction was performed with using the software GAPIT^43^. The models were run for each trial, and accuracy was measured from performing 5 fold cross validation in 10 replicates for each trial. Two models were run for each trait/trial. One model used a kinship matrix estimated 1 SNP for each of the associated genomic regions, while the other used 30,000 random SNPs for the estimation of the kinship matrix. All models included 10 MDS weights to account for population structure.

### Expression across tissues

We used the transcription data from the maize atlas^33^ for the following 11 tissues: 16 days after pollination embryo, 16 days after pollination endosperm, 6 days after silking primary root, tip of stage 2 leaf at V5 plant stage, base of stage 2 leaf at V5 plant stage, 13th leaf at V9 stage, 13th leaf at R2 stage, silk, anthers, Immature cob at V18 stage, 4th internode at V9 stage, and stem and shoot apical meristem at V4 stage. We used the standardized expression values, and estimated for each gene what tissue it was most expressed at. We then performed a chisquared test for each tissue to test if there were more genes expressed at the candidate or associating genes than expected under the null model of equal levels of the global expression pattern.

## Acknowledgments

This work was supported by SAGARPA (La Secretaría de Agricultura, Ganadería, Desarrollo Rural, Pesca y Alimentación), Mexico under the MasAgro (Sustainable Modernization of Traditional Agriculture) initiative, the NSF Grants #1238014 and #0922493, and the USDA-ARS. We would like to thank ICAMEX, BIDASem, and Dupont-Pioneer for assistance establishing phenotypic evaluation trials.

## Supplemental Figures

**Supplemental figure 1.**
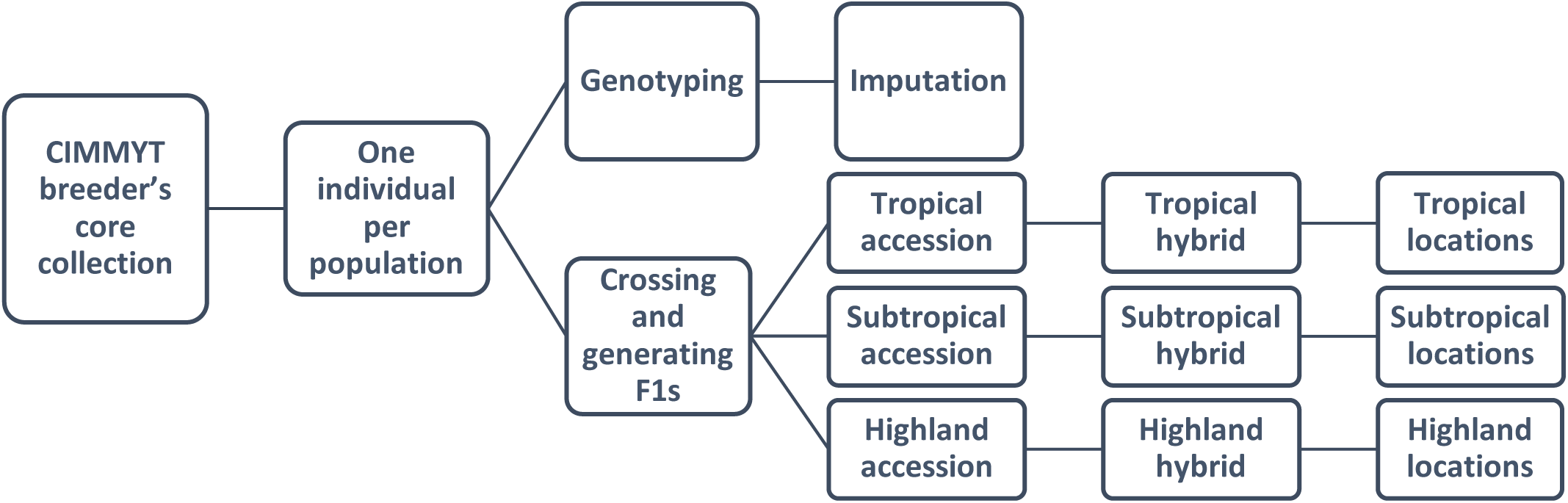
FOAM design with crossing and evaluation nested within adaptation.

**Supplemental figure 2.**
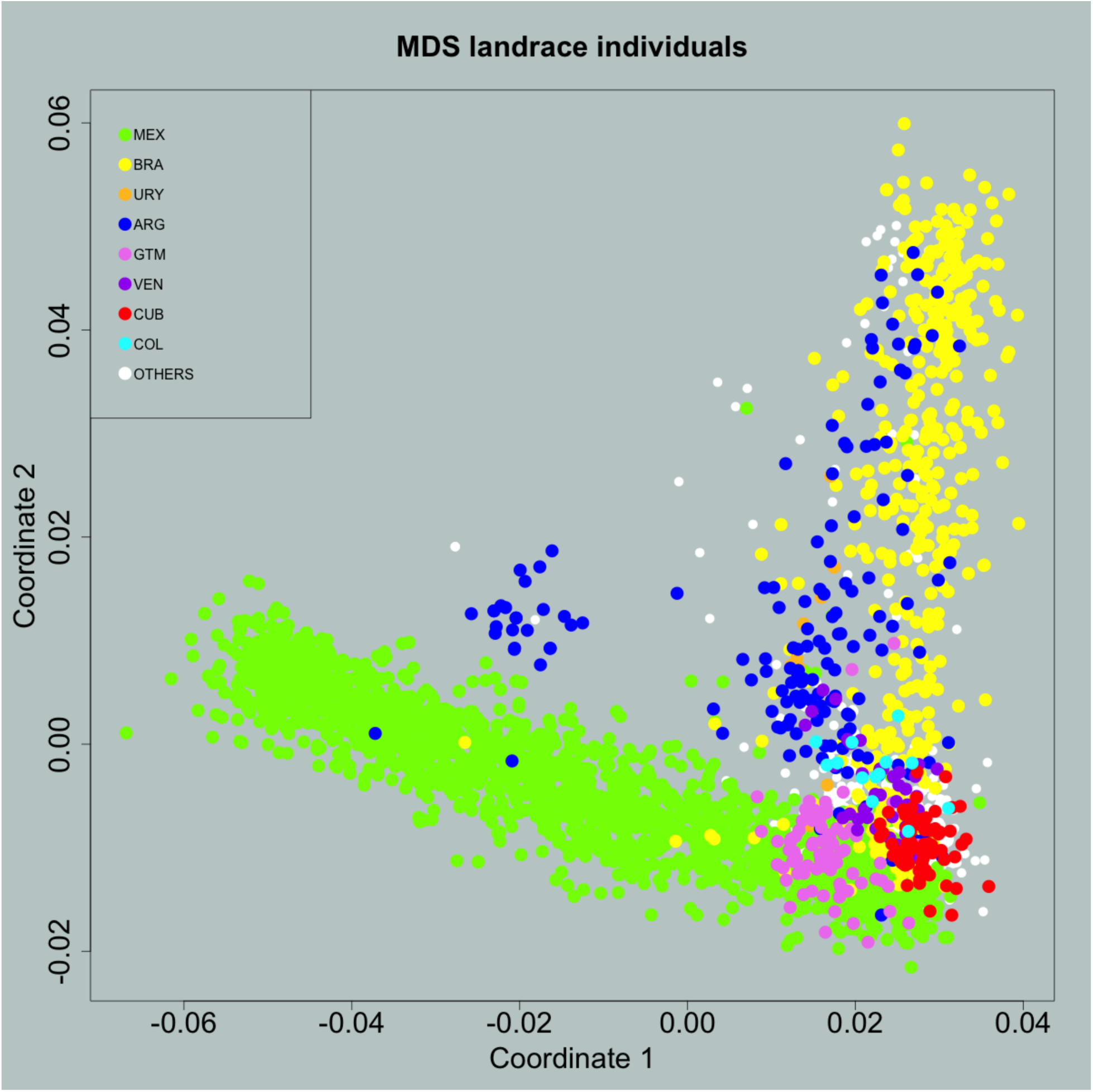
First 2 Principal Coordinates from Multidimensional scaling of the genetic distance among accessions

**Supplemental figure 3.**
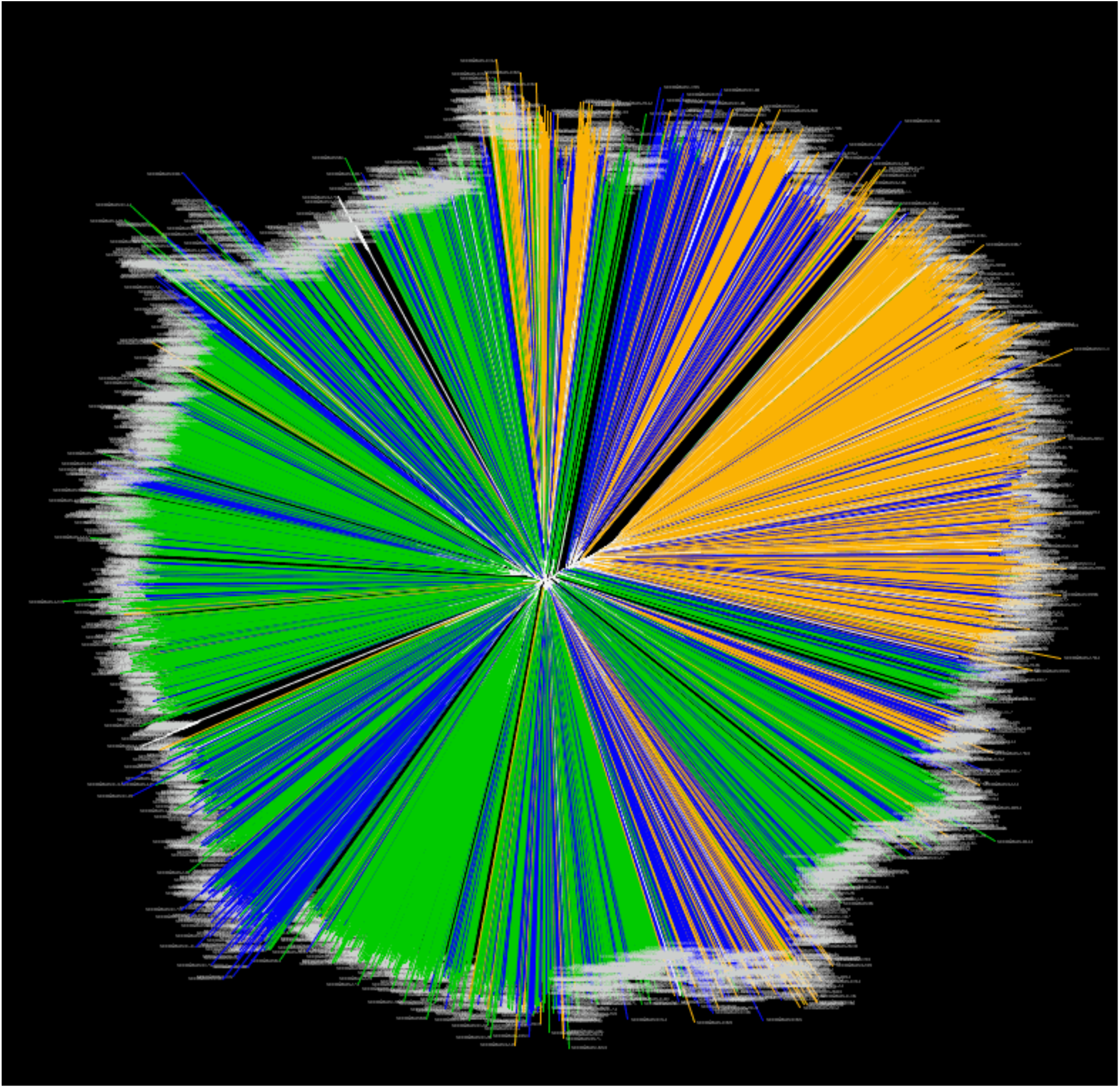
Neighbor-joining tree. Adaptation classes are colored green for low elevation, blue for mid elevation and orange for high elevation

**Supplemental figure 4.**
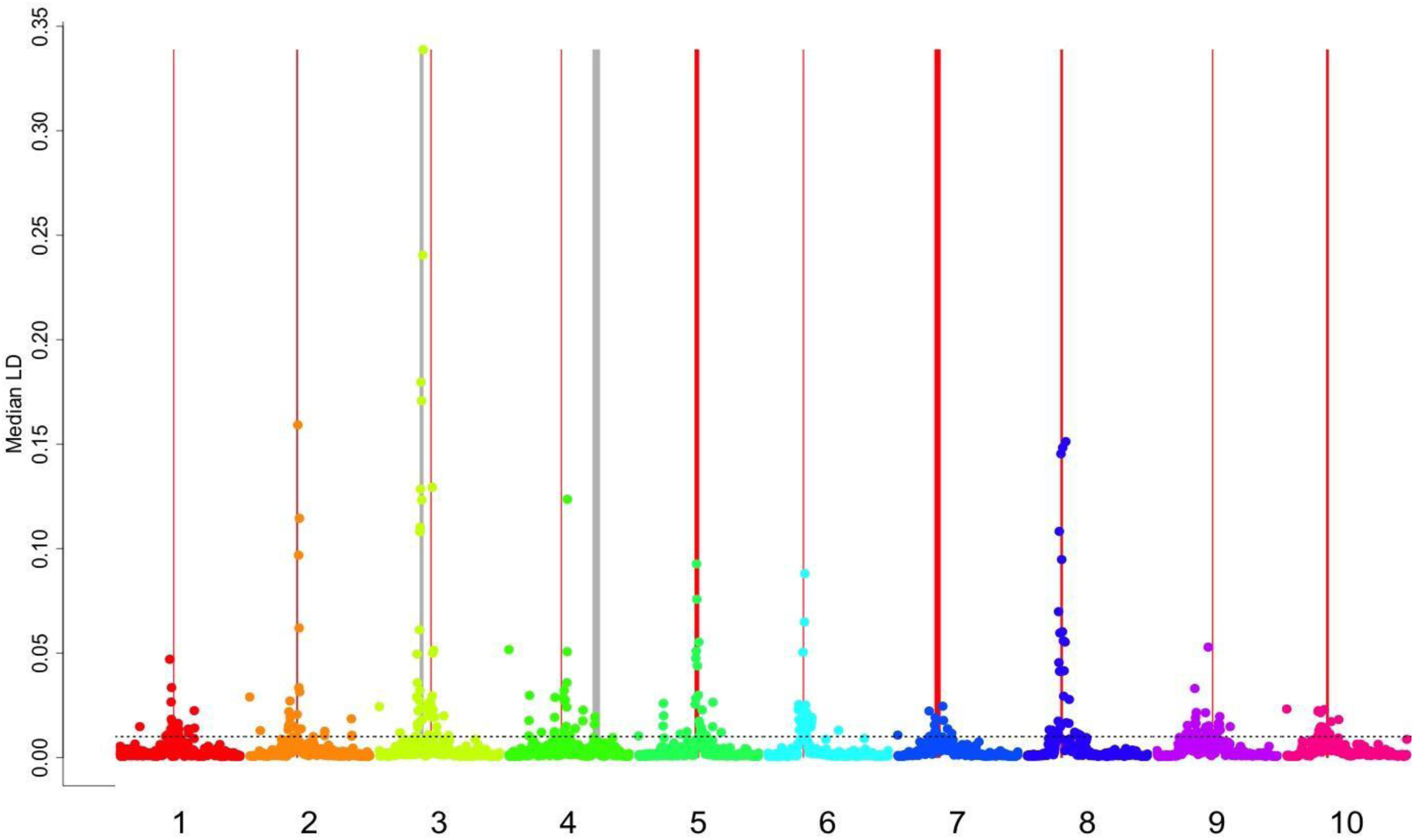
Genome wide view of the LD empirical threshold. Red shaded areas represent the centromeres, gray shaded areas represent inversions on chromosomes 3 and 4, and the dashed horizontal line represents the empirical LD threshold

**Supplemental figure 5.**
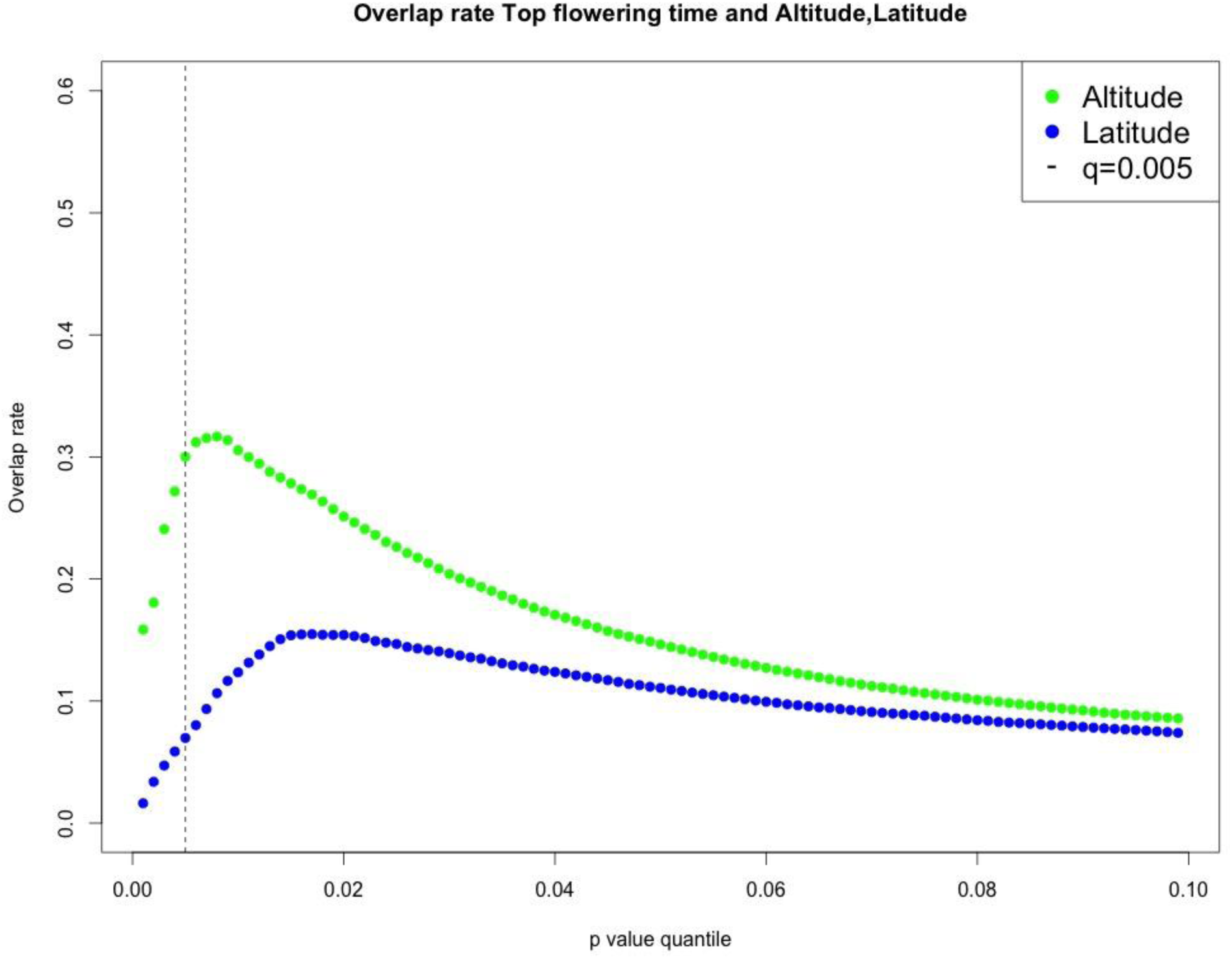
Overlap rate between the top associating SNPs with flowering time and altitude, latitude at various p-value thresholds.

**Supplemental figure 6.**
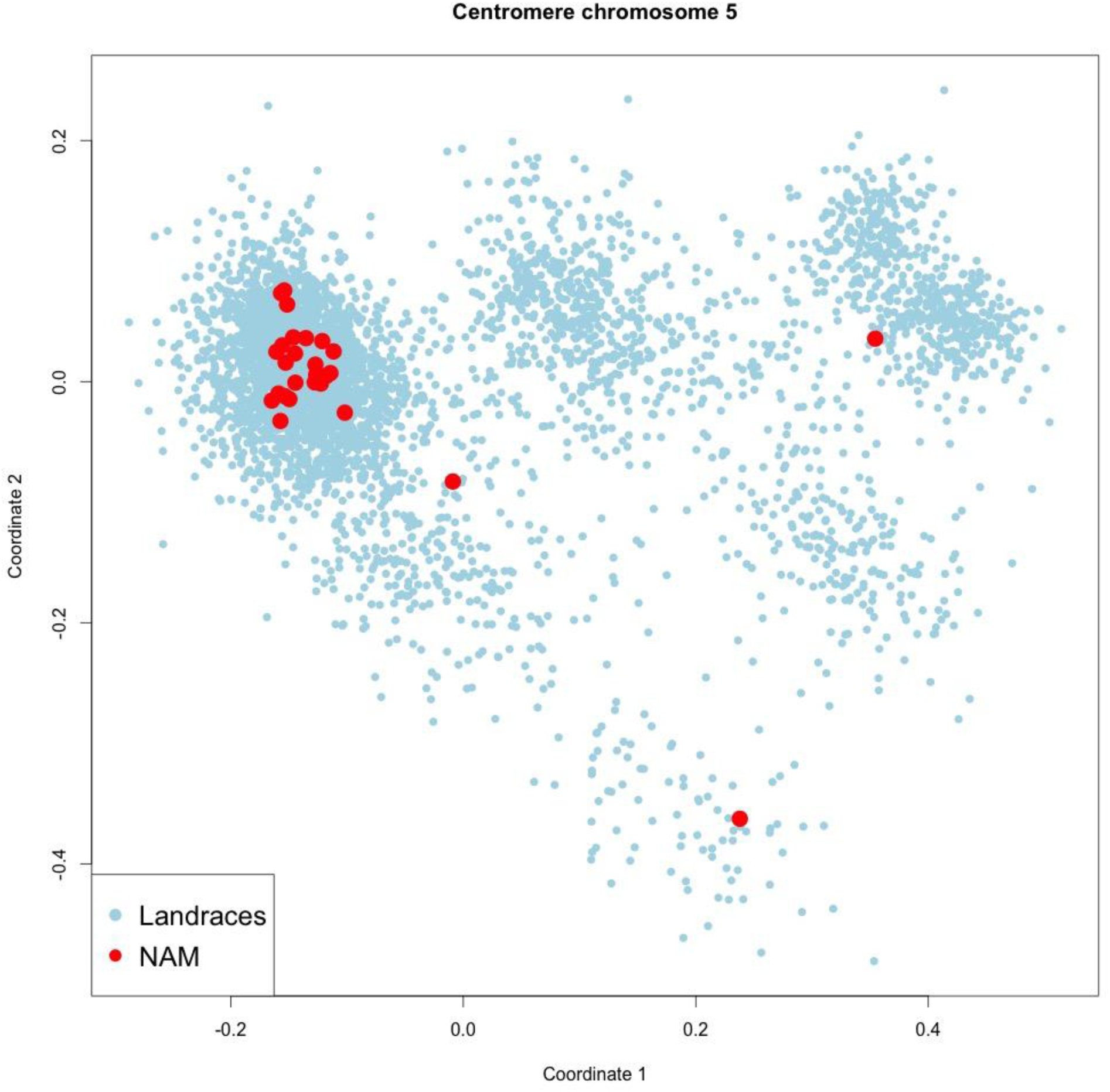
MDS of centromere of chromosome 5 for the FOAM landrace accessions and the NAM founders: Topright: Il14H. Bottom: P39. Middle: CML333

**Supplemental figure 7.**
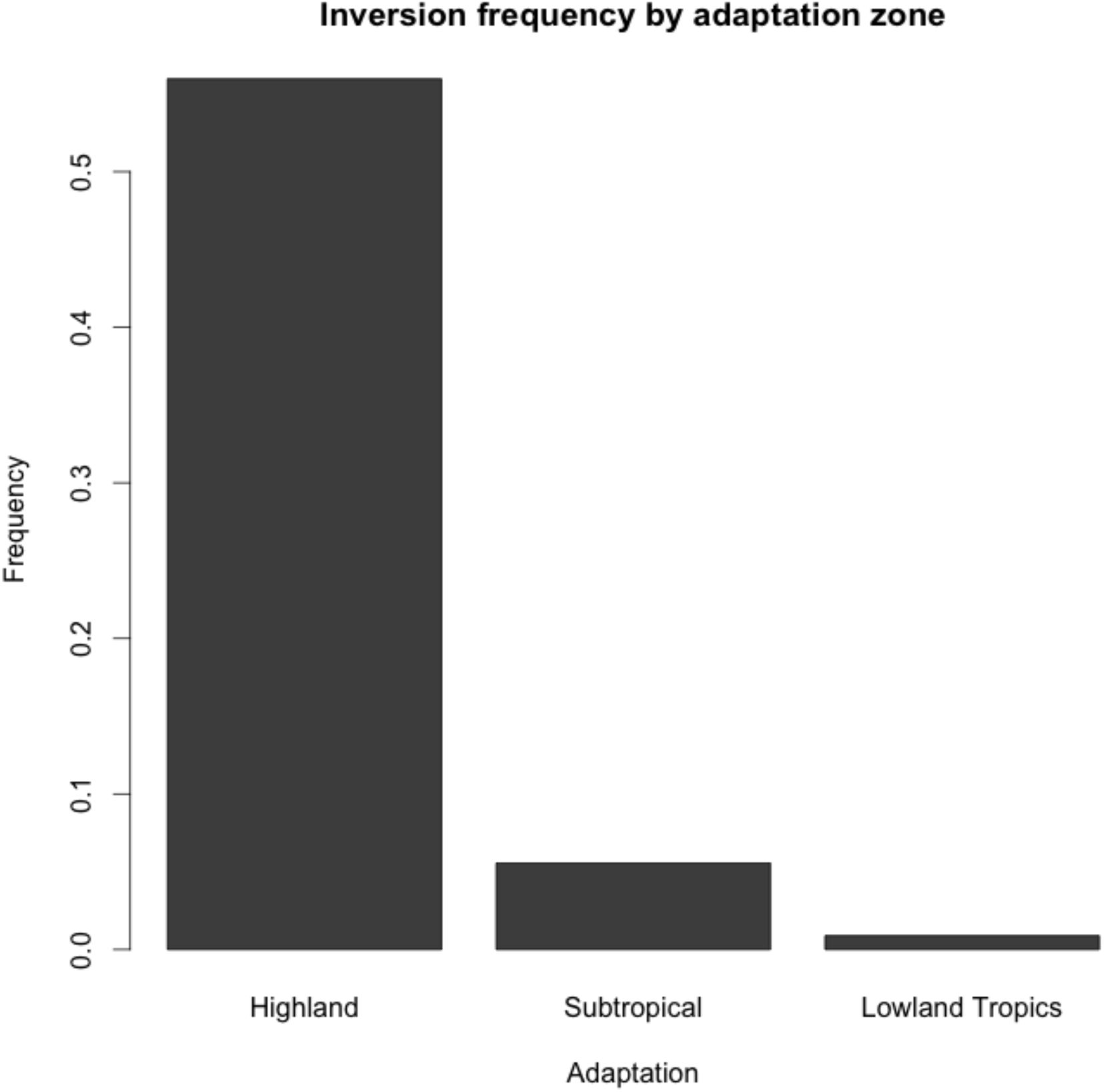
Frequency of INV4m according to accessions’ adaptation class

**Supplemental figure 8.**
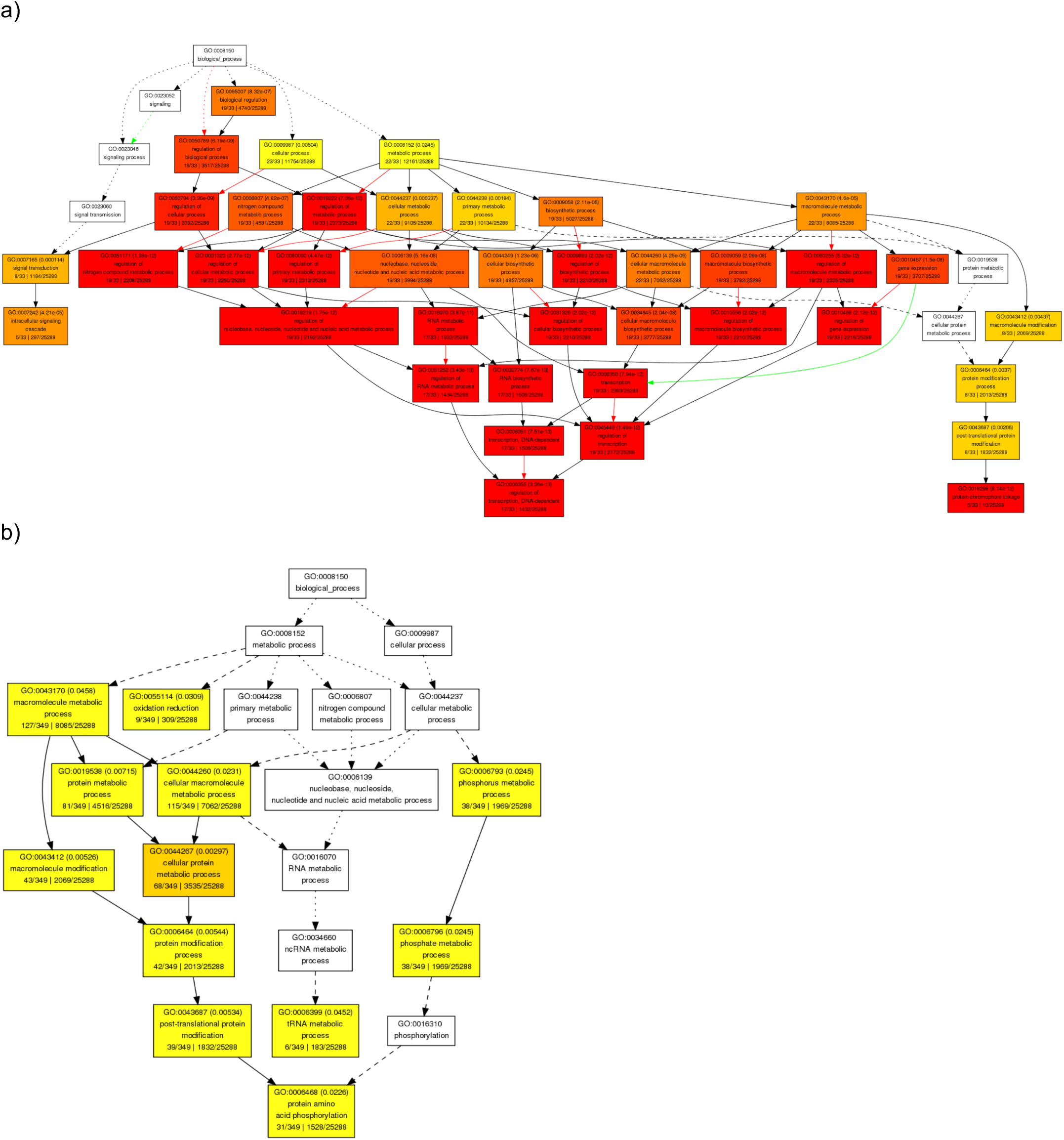
Ontology of genes for a) regulatory network genes and b) all associating gene

**Supplemental figure 9.**
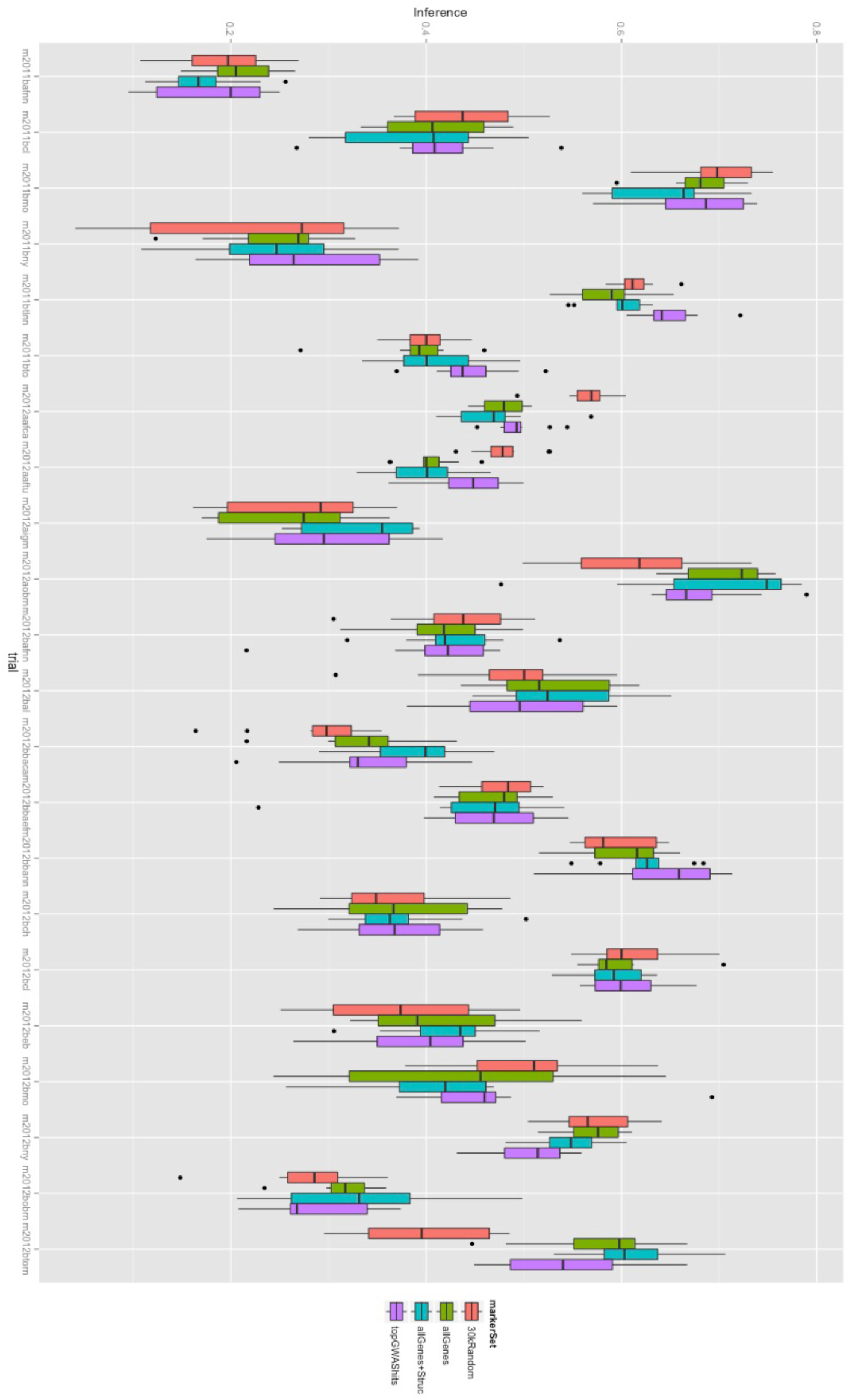
Genomic Prediction accuracy by trial

**Supplemental table 1.**
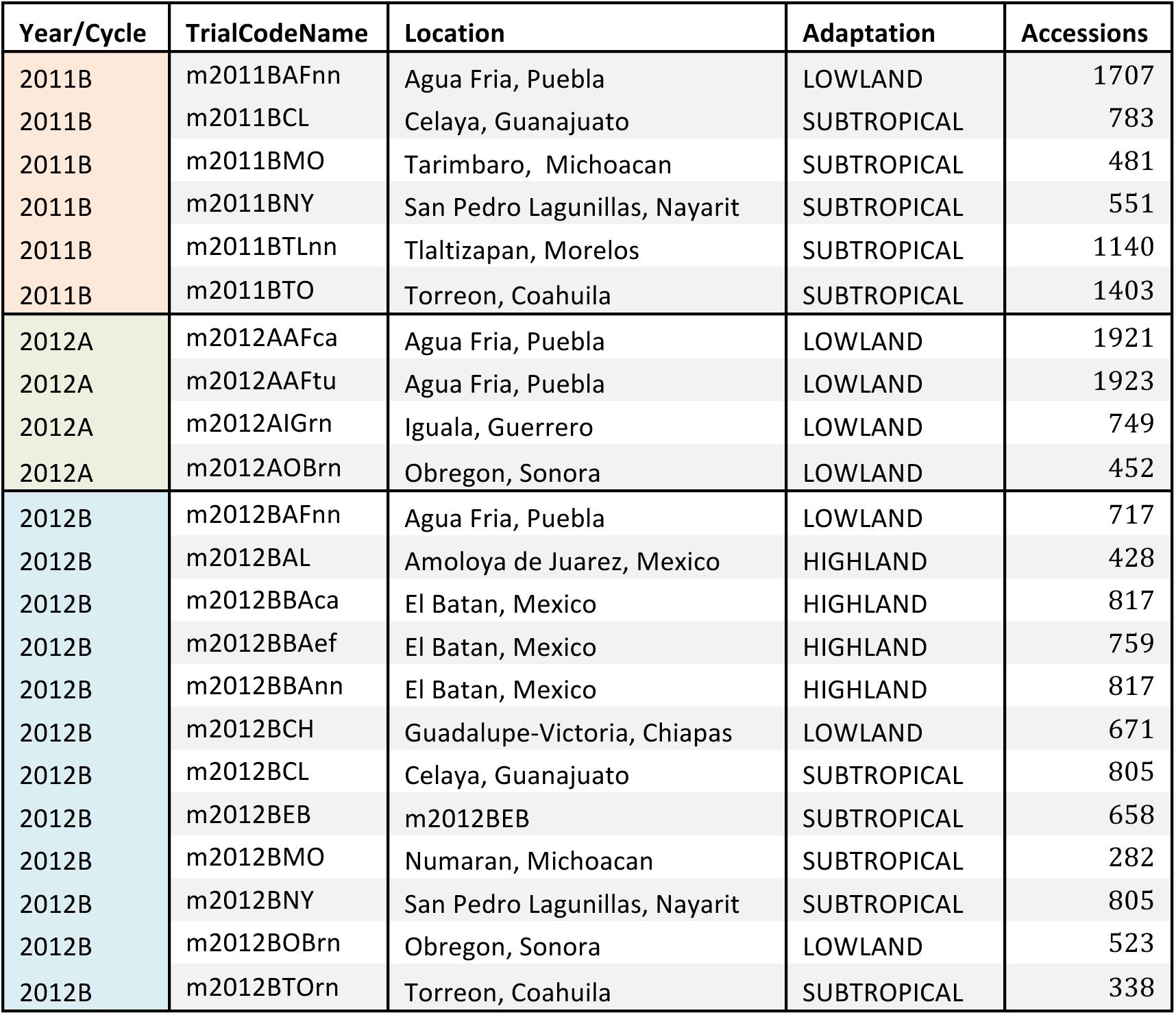
Trials location, adaptation class, and number of accessions planted

